## Extended data for "The mechanical anisotropy of adipose tissues regulates ovarian cancer invasion"

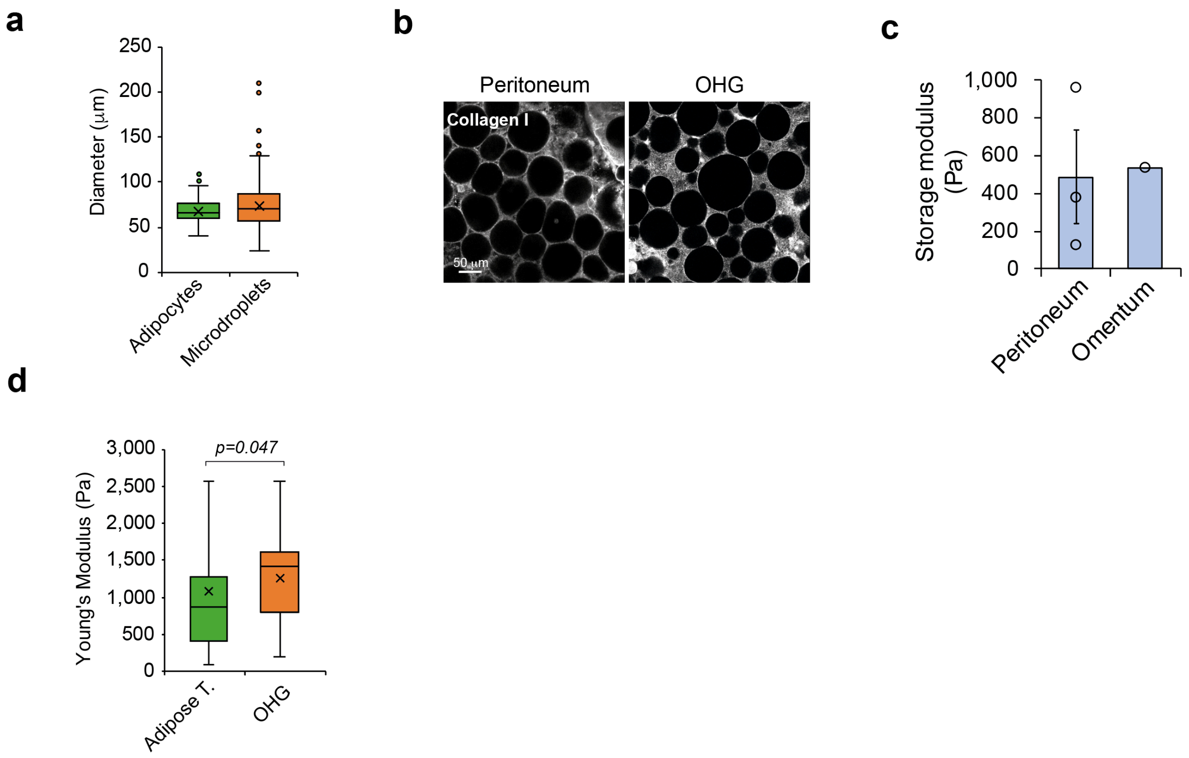


**Extended data Fig. 1: Comparison between human tissues and collagen-based OHG.**

**a**, Quantification of human peritoneal adipocyte and silicone oil droplet diameter (N = 170 cells/droplets). **b**, Collagen-I (grey) staining of human peritoneal adipose tissue and OHG. **c**, Storage modulus of human peritoneal and omental adipose tissues (N = 1-3). **d**, AFM quantification of the Young's modulus (N ≥ 208 measurements from ≥ 4 regions). Scale bars, 50 μm (**b**).

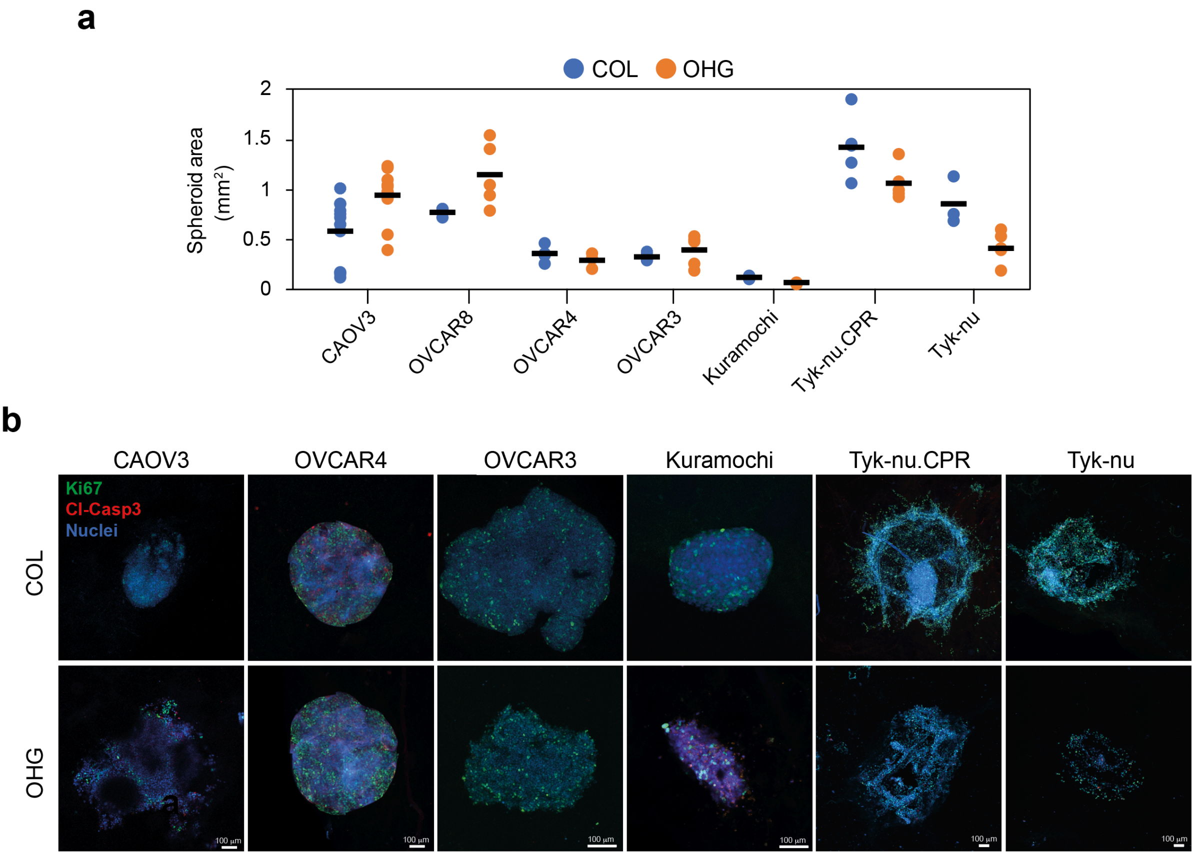


**Extended data Fig. 2: Ovarian cancer spheroid response to collagen and collagen-based OHG.**

**a**, Total spheroid area comparison of multiple ovarian cancer cell lines in collagen or OHGs (N ≥ 3 spheroids). **b**, Ki67 (green), cleaved-caspase 3 (red) and nuclei (blue) staining of spheroids embedded in collagen or OHG after 7 days. Scale bars, 100 μm (**b**).

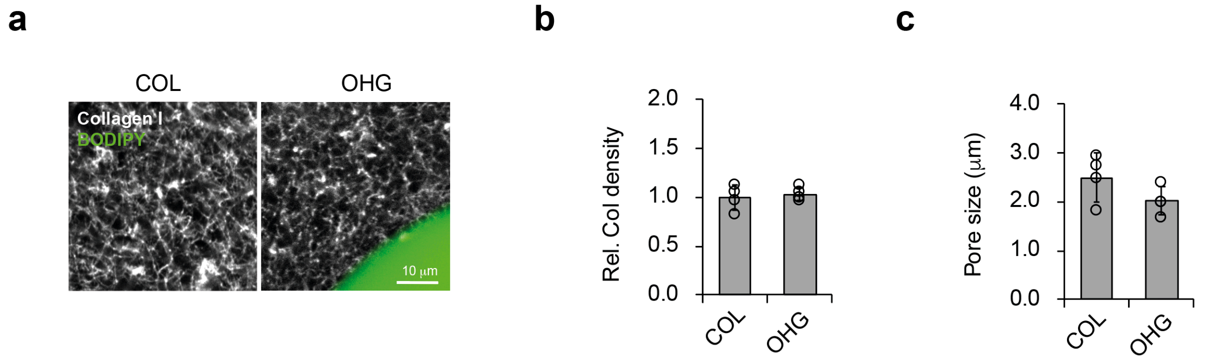


**Extended data Fig. 3: Invasion in OHG is enhanced compared to collagen gels.**

**a**, Collagen-I (grey) and BODIPY (green) staining of a collagen gel and an OHG. **c**, Quantification of collagen density in collagen gels and OHG (N = 4-5 gels). **d**, Quantification of pore size in collagen gels and OHG (N = 4 gels). For the data in **b** and **c**, a two-sided unpaired t-test was performed. Scale bar, 10 μm (**a**).

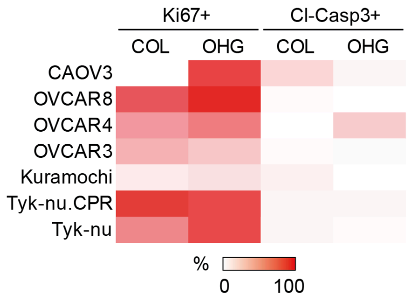


**Extended data Fig. 4: Ovarian Cancer cell proliferation and apoptosis in collagen and OHGs**

Average percentage of Ki67+ and cleaved-caspase 3+ cells in collagen or OHG (N > 75 cells).

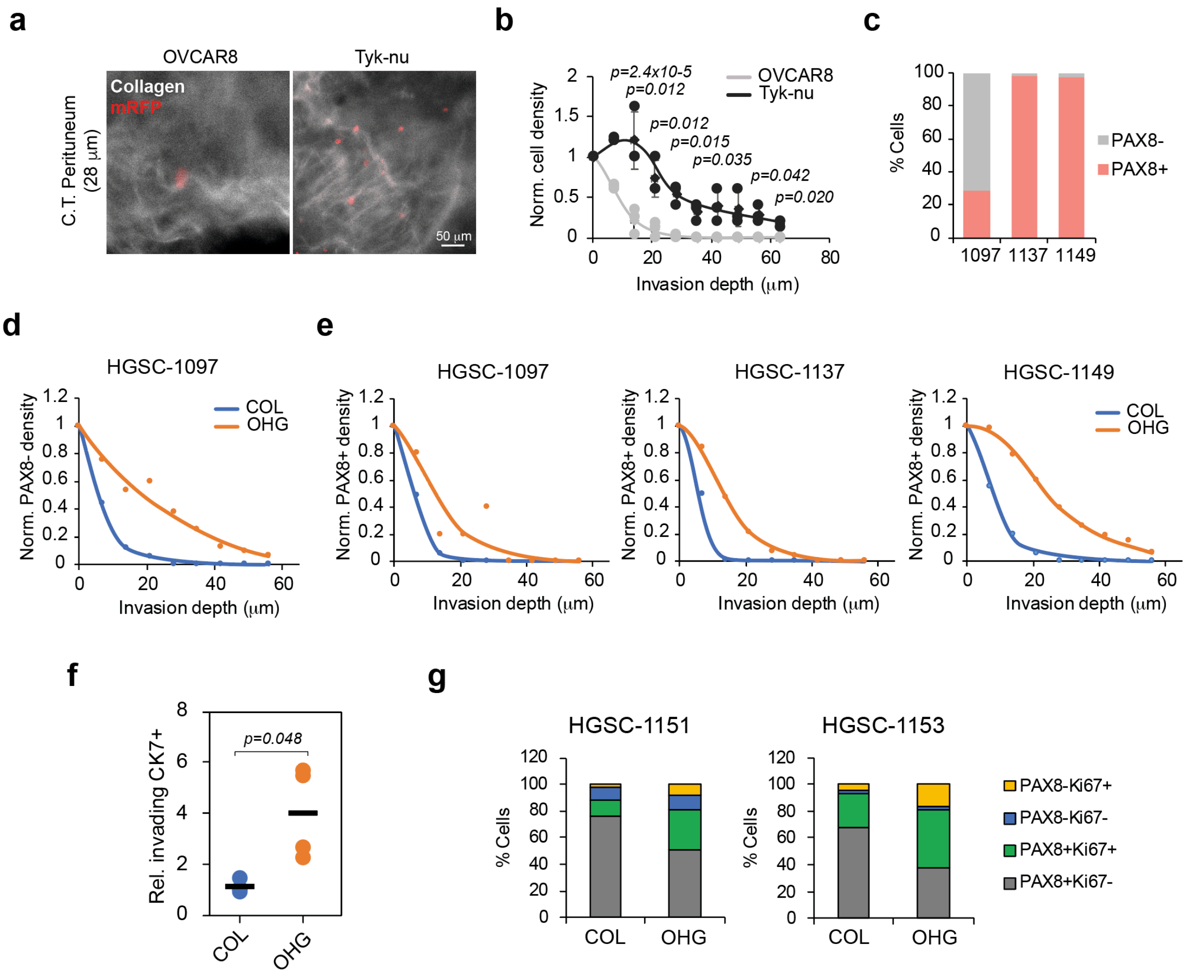


**Extended data Fig. 5: Invasion in OHG is enhanced compared to collagen gels.**

**a**, Collagen-I (grey) and mRFP (red) staining of mRFP-OVCAR8 or mRFP-Tyk-nu cells seeded on peritoneal connective tissue at 28 μm tissue depth. **b**, Quantification of OVCAR8 and Tyk-nu cell invasion depth into peritoneal connective tissue (N = 3 tissues from 1 donor). **c**, Percentage of PAX8- and PAX8+ cells in malignant ascites after 7 days in culture (N ≥ 246 cells). **d**, Quantification of PAX8- cell organotypic invasion depth after 7 days (N = 176 cells from one donor). **e**, Quantification of PAX8+ cell organotypic invasion depth after 7 days (N ≥ 70 cells in 3 donors). **f**, Quantification of invading single CK7+ cells relative to the number of spheroids from HGSC ascites cells embedded in collagen or OHG (N = 4 gels from 1 donor). **g**, Quantification of Ki67+ cells in PAX8+ and PAX8- cells of HGSC ascites cells embedded in collagen or OHG for 7 days (N ≥ 200 cells in two donors). For the data in **b** and **f** a two-sided unpaired t-test was performed. Scale bar, 50 μm (**a**).

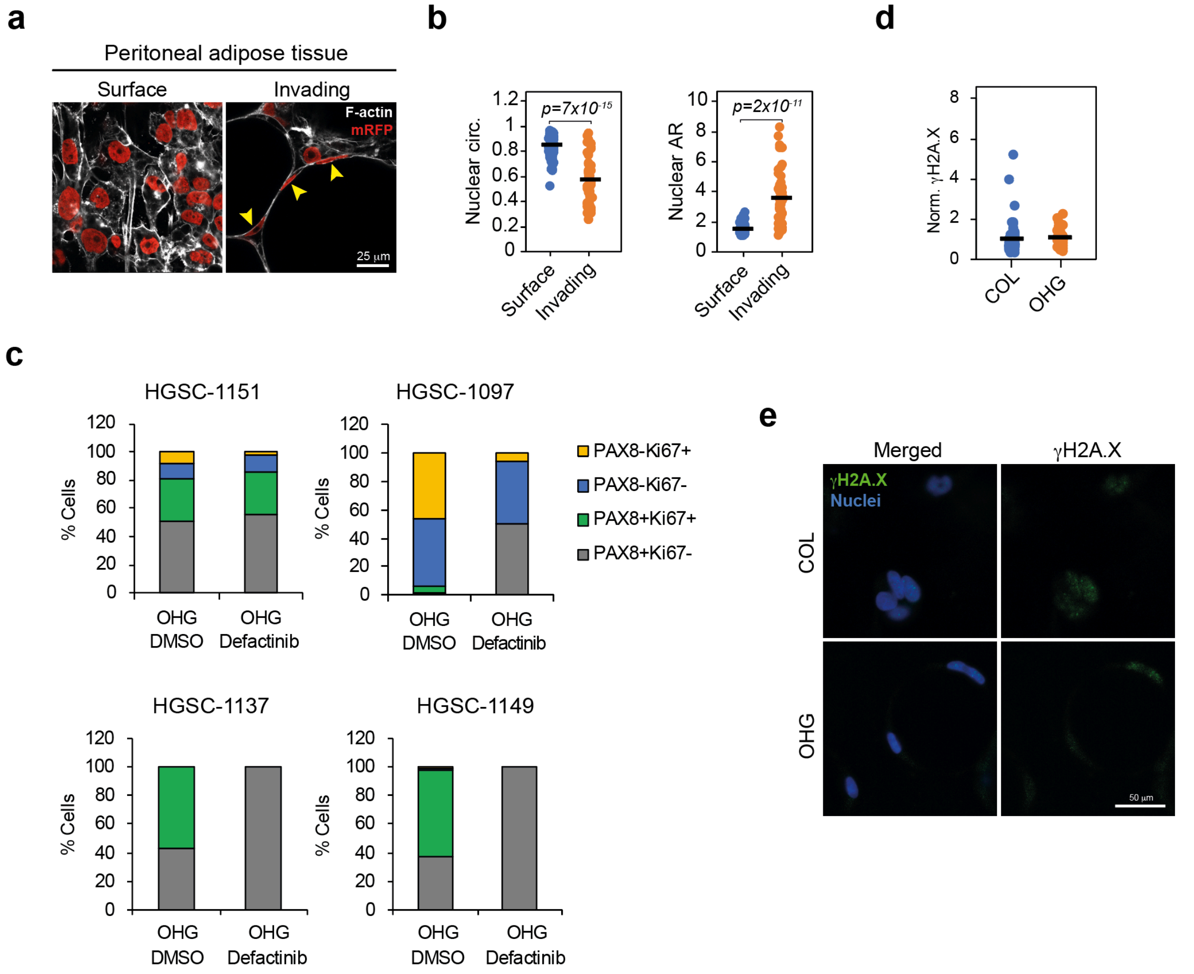


**Extended data Fig. 6: FAK-dependent cell invasion causes nuclear flattening but not DNA damage in ovarian cancer cells.**

**a**, F-actin (grey) and mRFP (red) staining of mRFP-OVCAR8 cells invading human peritoneal adipose tissue explants. **b**, Quantification of nuclear shape parameters in mRFP-OVCAR8 cells invading human peritoneal adipose tissue explants (N = 50-79 cells). **c**, Quantification of Ki67+ cells in PAX8+ and PAX8- cells of HGSC ascites cells embedded in collagen or OHG for 7 days (N ≥ 43 cells in 4 donors). **d**, Normalised intensity of γH2A.X immunofluorescence staining in OVCAR8 cells in collagen or OHG (N ≥ 38 nuclei). **e**, γH2A.X (green) and nuclei (blue) staining of OVCAR8 cells in collagen or OHG. For the data in **b** and **d** a two-sided unpaired t-test was performed. Scale bars, 25 μm (**a**) and 5o μm (**e**).

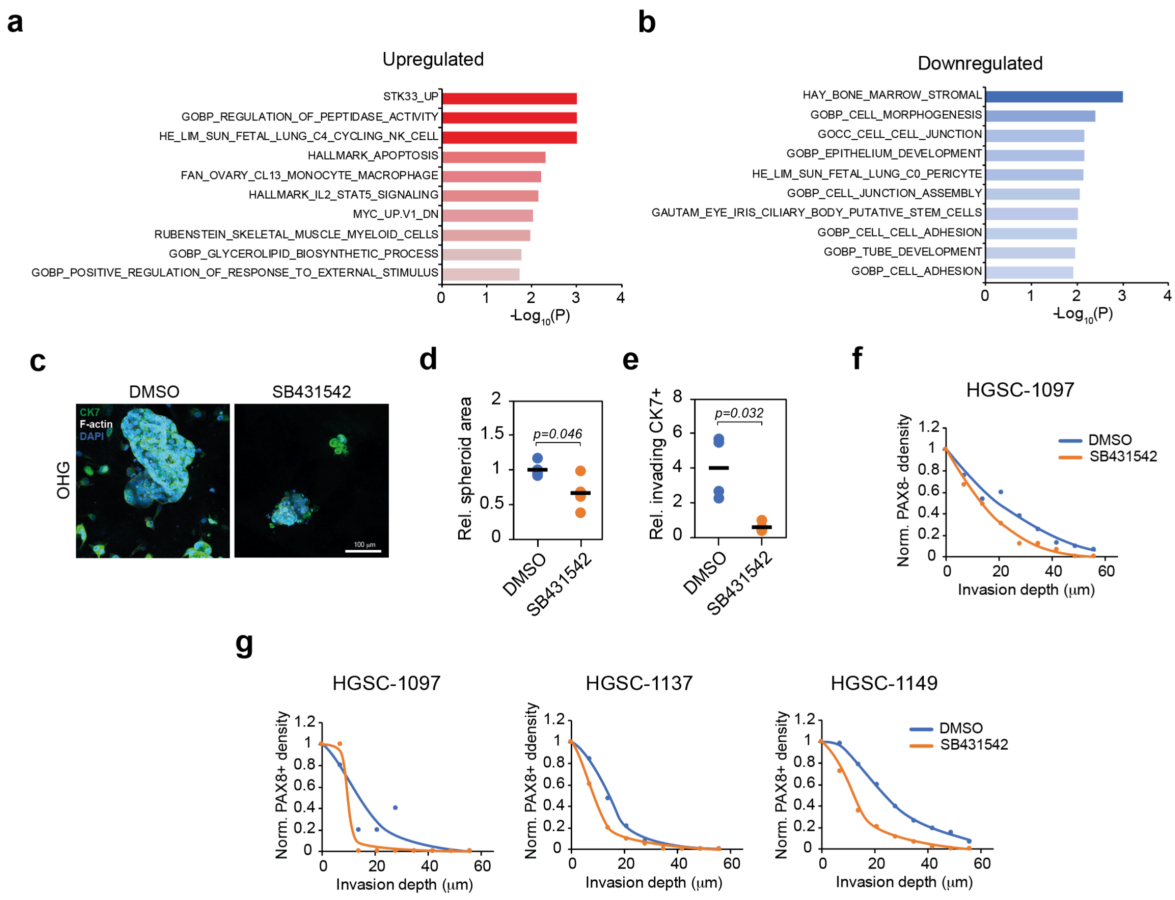


**Extended data Fig. 7: TGFβ signalling promotes invasion in OHG.**

**a**, The ten most enriched gene sets from the most significantly upregulated genes in OHG-invasive ovarian cancer cell lines compared to non-invasive lines. **b**, Ten most enriched gene sets from the significantly downregulated genes in OHG-invasive ovarian cancer cell lines compared to non-invasive lines. **c**, CK7 (green), F-actin (grey), and DAPI (blue) staining of HGSC ascites cells embedded in OHG. **d**, Quantification of spheroid area of HGSC ascites cells embedded in OHG (N = 4 gels from 1 donor). **e**, Quantification of invading single CK7+ cells relative to the number of spheroids from HGSC ascites cells embedded in OHG (N = 4 gels from 1 donor). **f**, Quantification of PAX8- cell organotypic invasion depth after 7 days (N = 212 cells from one donor). **g**, Quantification of PAX8+ cell organotypic invasion depth after 7 days (N ≥ 125 cells in 3 donors). For the data in **d** and **e** a two-sided unpaired t-test was performed. Scale bar, 100 μm (**c**).

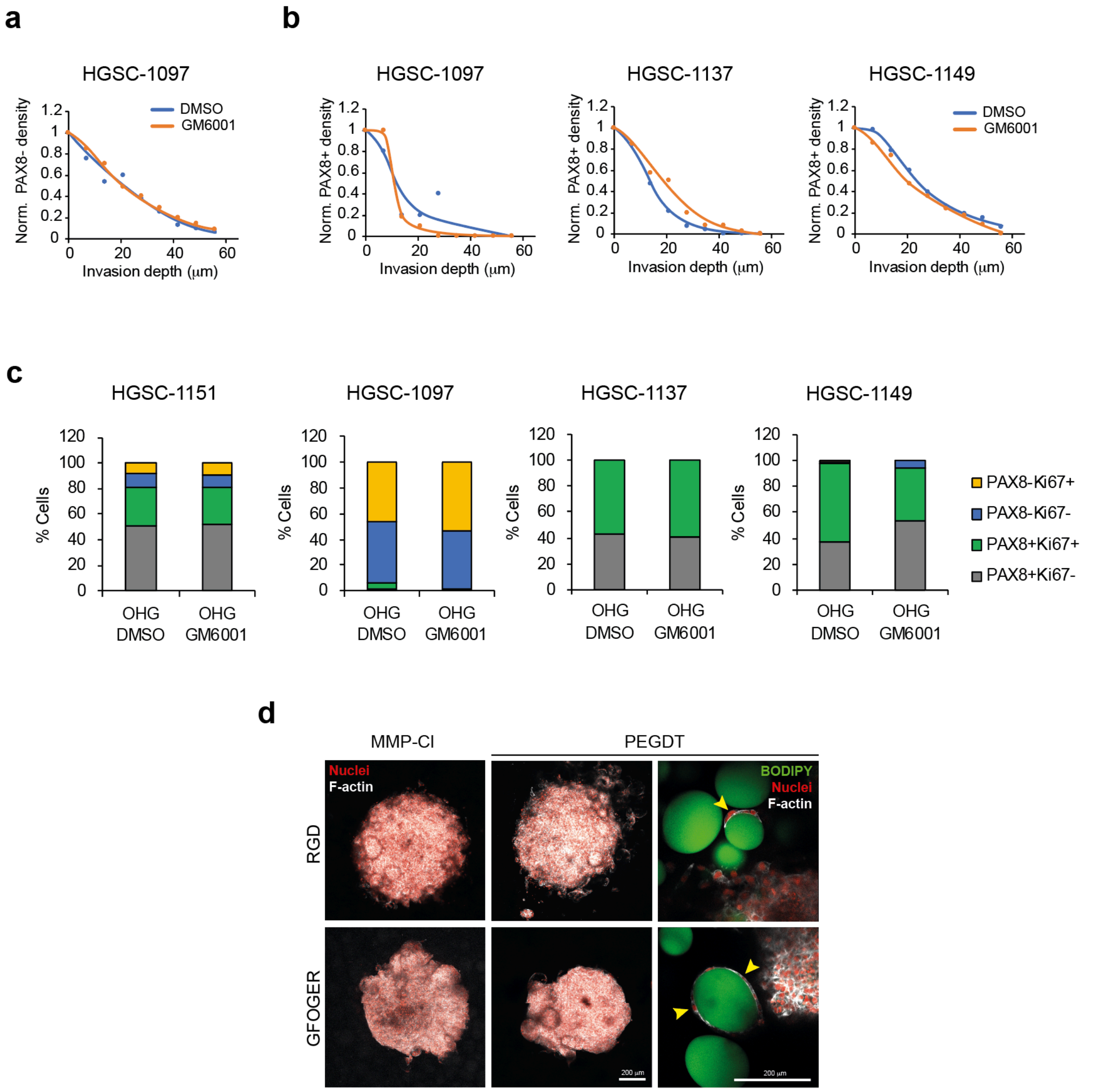


**Extended data Fig. 8: Cell invasion into adipose tissue is MMP-independent.**

**a**, Quantification of PAX8- cell organotypic invasion depth after 7 days (N = 504 cells from one donor). **b**, Quantification of PAX8+ cell organotypic invasion depth after 7 days (N ≥ 38 cells in 3 donors). **c**, Quantification of Ki67+ cells in PAX8+ and PAX8- cells of HGSC ascites cells embedded in OHG for 7 days (N ≥ 54 cells in 4 donors). **d**, BODIPY (green), nuclei (red) and F-actin (grey) staining of OVCAR8 spheroids in MMP-Cleavable or non-cleavable (PEGDT) HA-NB OHG. Scale bars, 200 μm (**d**).

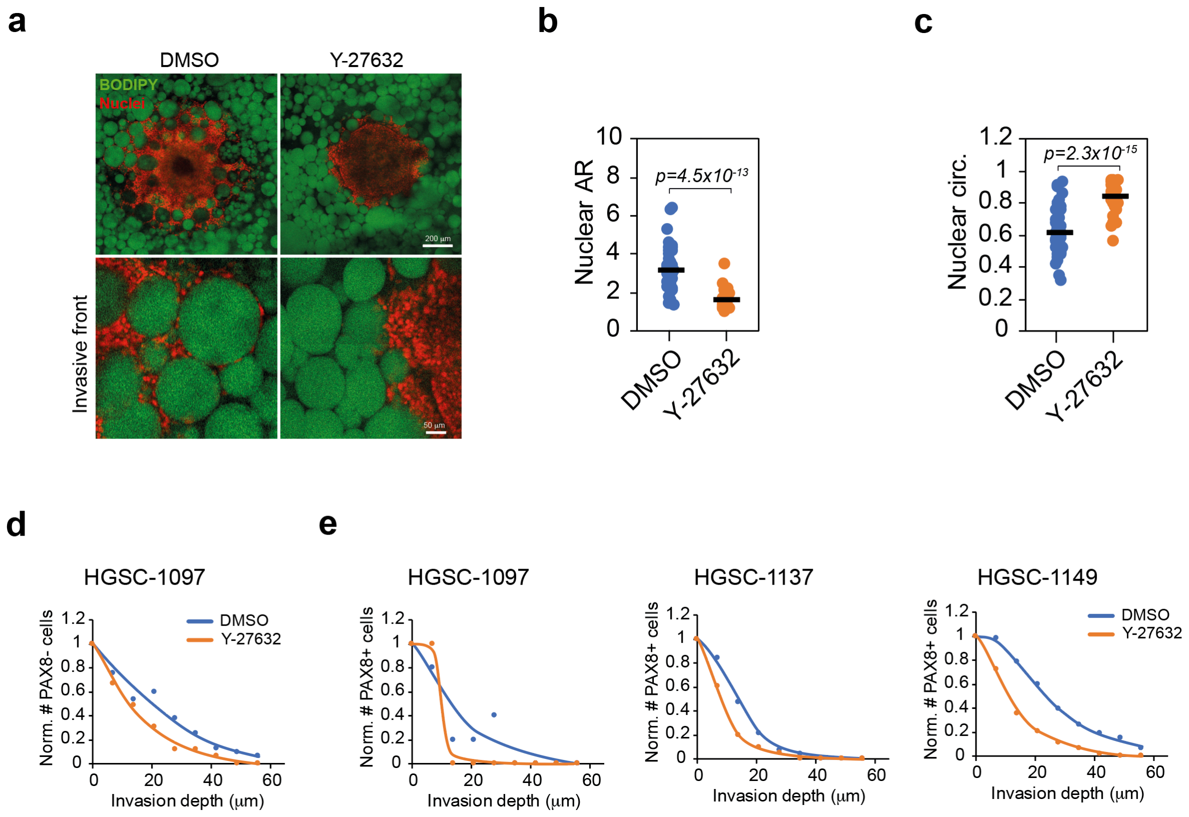


**Extended data Fig. 9: Cell invasion into adipose tissue is ROCK-dependent.**

**a**, BODIPY (green) and nuclei (red) staining of OVCAR8 cells in collagen-based OHG. **b**, Quantification of nuclear aspect ratio of OVCAR8 cells (N = 50 cells). **c**, Quantification of nuclear circularity of OVCAR8 cells (N = 50 cells). **d**, Quantification of PAX8- cell organotypic invasion depth after 7 days (N = 158 cells from one donor). **e**, Quantification of PAX8+ cell organotypic invasion depth after 7 days (N ≥ 79 cells in 3 donors). For the data in **b** and **c** a two-sided unpaired t-test was performed. Scale bars, 50 μm (bottom **a**), 200 μm (top **a**).

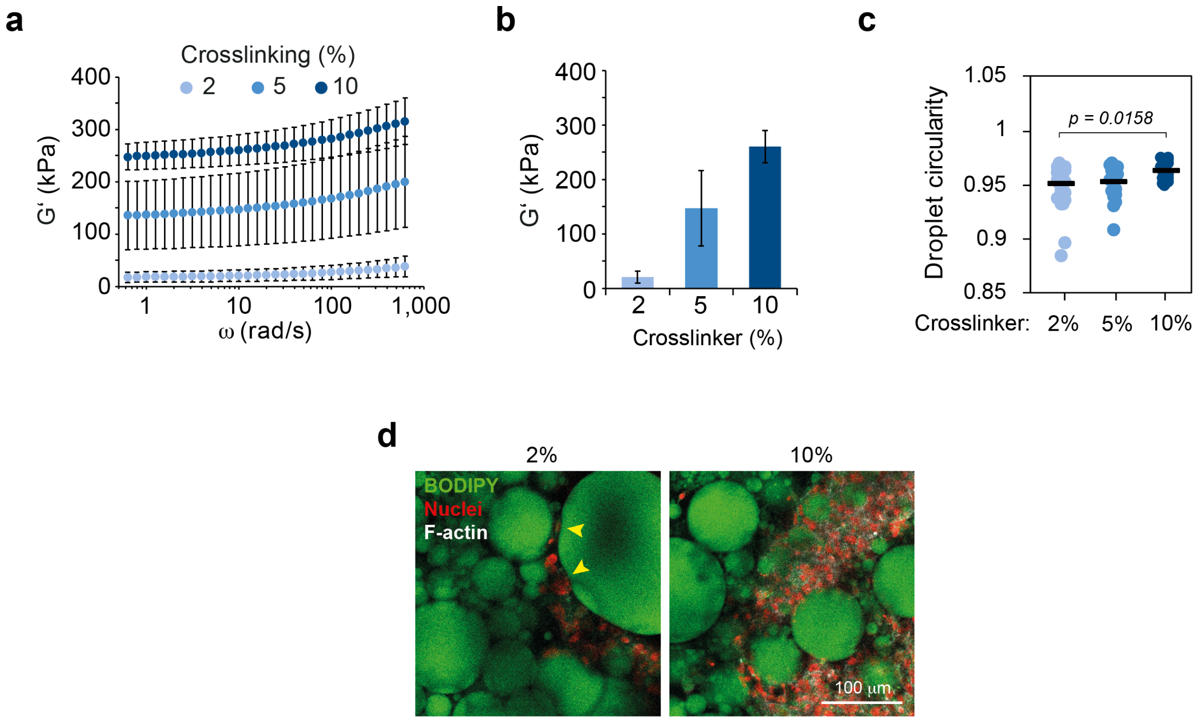


**Extended data Fig. 10: Cell-induced deformation is prevented in stiff PDMS microbeads.**

**a**, **b**, Frequency sweeps (**a**) and shear storage moduli (at 10 rad/s) (**b**) of Sylgard 184 PDMS cured with increasing crosslinker concentrations (N = 3). **c**, Quantification of microbead circularity of microbeads in contact with OVCAR8 cells (N = 25 microbeads). **d**, BODIPY (green), F-actin (grey), and nuclei (red) staining of OVCAR8 spheroids embedded in PDMS OHGs crosslinked with collagen. Yellow arrowheads indicate deformations caused by OVCAR8 cells. One-way analysis of variance (ANOVA) with Tukey's correction for multiple comparisons was performed for the data in **c**. Scale bar, 100 μm (**d**).

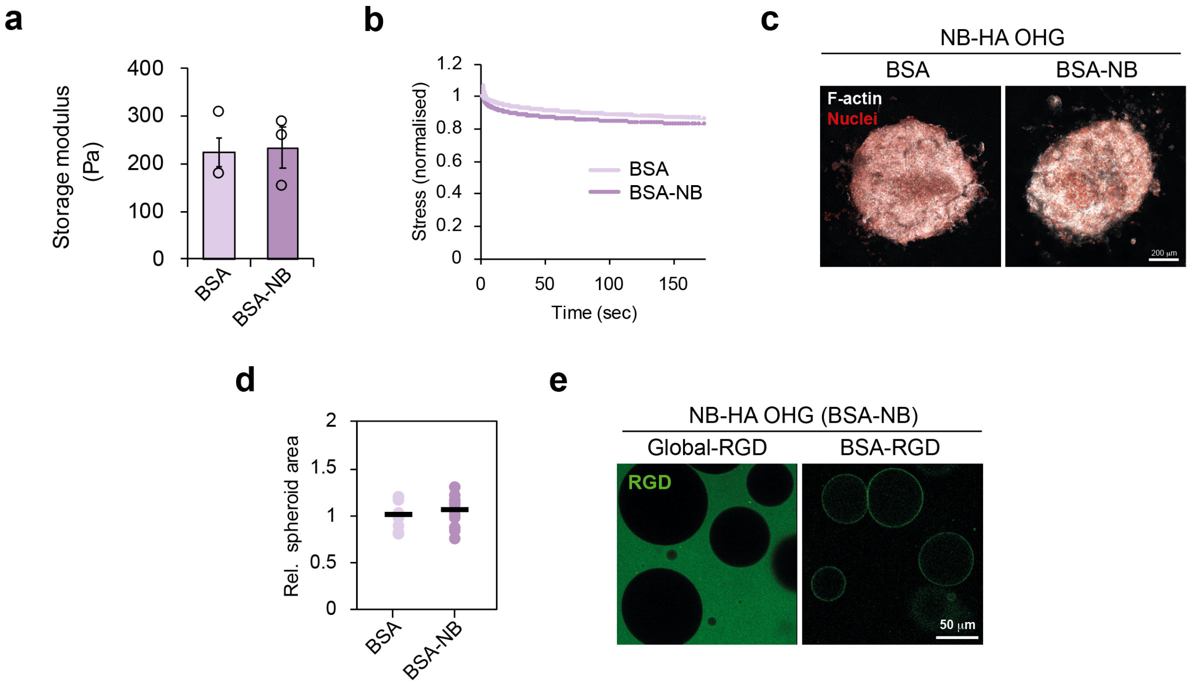


**Extended data Fig. 11: Mechanical characterisation and RGD localisation in NB-HA OHGs.**

**a**, Quantification of storage modulus of HA-NB OHG formed with emulsions stabilised with BSA or BSA-NB nanosheets and crosslinked with PEGDT (N = 3 gels). **b**, Normalised stress-relaxation curves of HA-NB OHG formed with emulsions stabilised with BSA or BSA-NB nanosheets and crosslinked with PEGDT (N = 3 gels). **c**, F-actin (grey) and nuclei (red) staining of OVCAR8 spheroids after 7 d culture embedded in HA-NB OHG containing BSA or BSA-NB (crosslinked) microdroplets. **d**, Quantification of relative OVCAR8 spheroid area after culture in BSA or BSA-NB NB-HA OHG (N = 11-12 spheroids). **e**, RGD-FITC (green) in HA-NB OHG with controlled RGD localisation. For the data in **a** and **d** a two-sided unpaired t-test was performed. Scale bars, 50 µm (**e**), 200 µm (**c**).

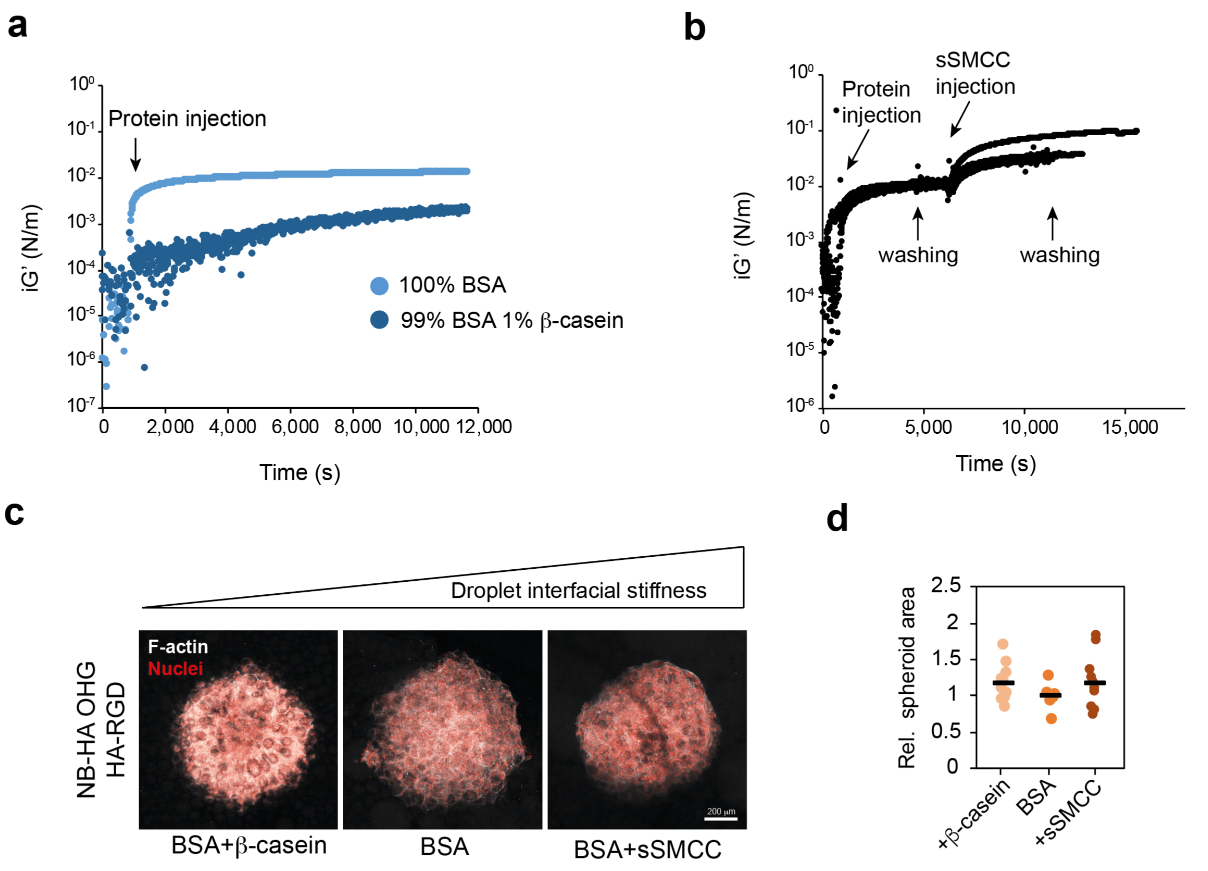


**Extended data Fig. 12: Interfacial protein nanosheet mechanics.**

**a**, Evolution of the interfacial shear storage modulus of 1 mg/ml BSA and BSA combined with β-casein protein nanosheets formed over NOVEC 7500 oil (0.1 Hz, 1.0x $10^{-4}$ rad). **b**, Evolution of the interfacial shear storage modulus of 1 mg/ml BSA before and after crosslinking with 2 mg/ml sulfo-SMCC. **c**, F-actin (grey) and nuclei (red) of OVCAR8 spheroids in HA-NB OHGs with BSA microdroplet (i.e. RGD presented in the HA phase but not on the microdroplet surface) and varying protein nanosheet interfacial mechanics. **d**, Relative OVCAR8 spheroid area in HA-NB OHGs with HA-presenting RGD and varying protein nanosheet interfacial mechanics (N = 9-11 spheroids). One-way analysis of variance (ANOVA) with Tukey's correction for multiple comparisons was performed for the data in **d**. Scale bar, 200 μm (**c**).

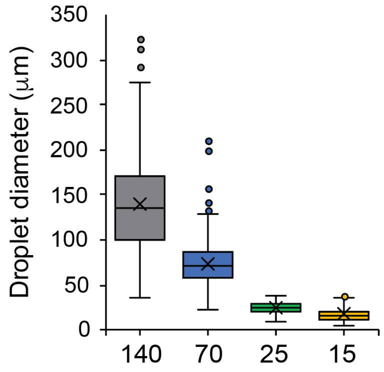


**Extended data Fig. 13: Diameter of oil microdroplets used in OHGs.**

Quantification of droplet diameter of emulsions (N = 170 droplets).

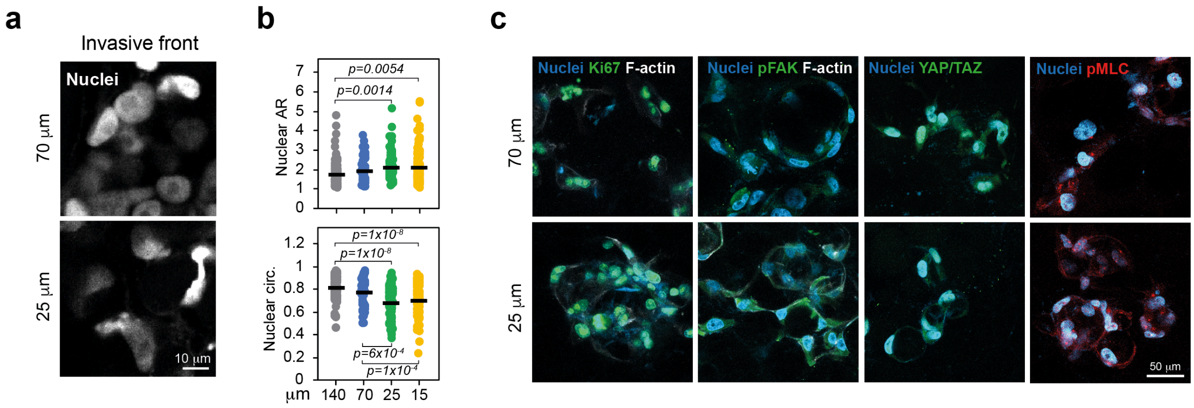


**Extended data Fig. 14: Droplet size in OHG regulates cell invasion.**

**a**, Nuclei (grey) staining of OVCAR8 cells at the invasive front of spheroids embedded in OHG of distinct droplet size. **b**, Quantification of nuclear shape descriptors of OVCAR8 cells from the invasive front of spheroids embedded in OHG of varying droplet size. **c**, Nuclei (blue), F-actin (grey), Ki67+, pFAK and pMLC (green, left to right) and pMLC (red) staining of OVCAR8 cells embedded in OHG of 70 or 25 µm diameter droplets. One-way analysis of variance (ANOVA) with Tukey's correction for multiple comparisons was performed for the data in **a**. Scale bars, 10 µm (**a**), 50 µm (**c**).

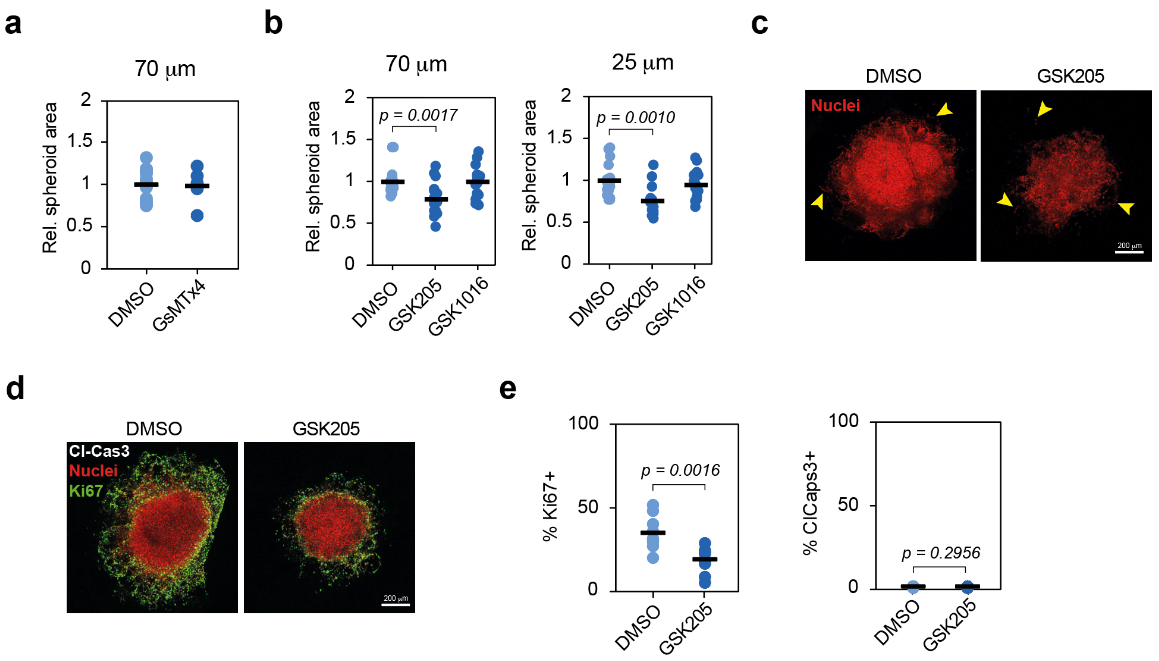


**Extended data Fig. 15: TRPV4 activity regulates cell proliferation.**

**a**, Quantification of OVCAR8 spheroid area in OHG formed with 70 µm droplets upon inhibition with Piezo1 inhibitor GsMTx4 (N = 5-16 spheroids). **b**, OVCAR8 spheroid area in OHG formed with 70 or 25 µm droplets upon treatment with the TRPV4 inhibitor GSK205 or agonist GSK1016790A (N ≥ 15 spheroids). **c**, Nuclei (red) of OVCAR8 cells embedded in OHG. Yellow arrowheads indicate cells at the invasive front. **d**, Cleaved Caspase 3 (grey), nuclei (red) and Ki67 (green) staining of OVCAR8 spheroids embedded in 70 µm droplet OHG for 7 days. **e**, Quantification of Ki67+ and Cleaved caspase 3+ cells from (**d**) (N ≥ 11,779 cells from 9 spheroids). For the data in **a** and **e**, a two-sided unpaired t-test was performed. One-way analysis of variance (ANOVA) with Tukey's correction for multiple comparisons was performed for the data in **b**. Scale bar, 200 µm (**c** and **d**).

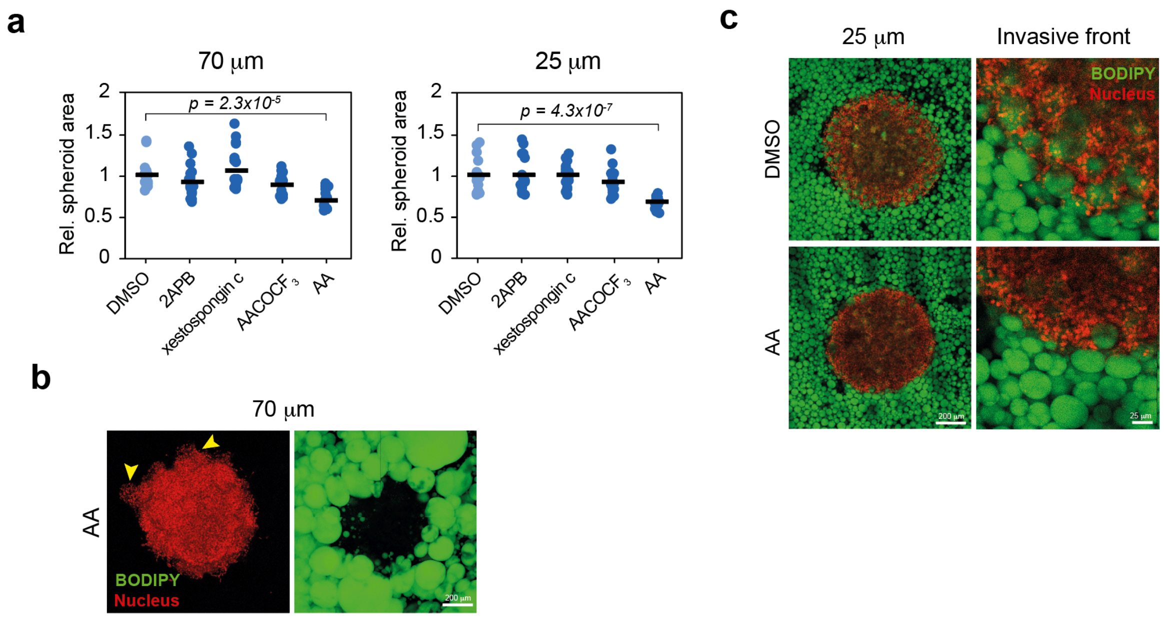


**Extended data Fig. 16: Inhibition of the cPLA2-Arachidonic acid-InsP3Rs pathway does not impair cell invasion.**

**a**, Quantification of OVCAR8 spheroid area in OHG formed with 70 or 25 μm droplets with perturbed mechanical confinement signalling (N ≥ 16 spheroids). **b**, BODIP (green) and nuclei (red) staining of OVCAR8 cells treated with arachidonic acid and embedded in 70 μm OHG. Yellow arrowheads indicate collectively-invading cells. **c**, BODIPY (green) and nuclei (red) of OVCAR8 cells embedded in 25 μm OHG. One-way analysis of variance (ANOVA) with Tukey's correction for multiple comparisons was performed for the data in **a**. Scale bars, 200 μm (**b**, left **c**), 25 μm (right **c**).
